# Revealing the transcription factor regulatory context of human specific cortical development using single-cell multi-omics

**DOI:** 10.1101/2021.03.19.436193

**Authors:** Yan Wu, Blue Lake, Brandon Sos, Song Chen, Thu E. Duong, Yun C. Yung, Weixiu Dong, Siddarth Limaye, Jerold Chun, Kun Zhang

## Abstract

Human behaviors are at least partially driven by genomic regions that influence human-specific neurodevelopment. This includes genomic regions undergoing human specific sequence acceleration (Human Accelerated Regions or HARs) and regions showing human-specific enhancer activity (Human Gained Enhancers or HGEs) not present in other primates. However, prior studies on HAR/HGE activities involved mixtures of brain cell types and focused only on putative downstream target genes. Here, we directly measured cell type specific HAR/HGE activity in the developing fetal human brain using two independent single-cell chromatin accessibility datasets with matching single-cell gene expression data. Transcription factor (TF) motif analyses identified upstream TFs binding to HARs/HGEs and identified LHX2, a key regulator of forebrain development, as an active HGE regulator in neuronal progenitors. We integrated our TF motif analyses with published chromatin interaction maps to build detailed regulatory networks where TFs are linked to downstream genes via HARs/HGEs. Through these networks, we identified a potential regulatory role for NFIC in human neuronal progenitor networks via modulating the Notch signaling and cell adhesion pathways. Therefore, by using a single cell multi-omics approach, we were able to capture both the upstream and downstream regulatory context of HARs/HGEs, which may provide a more comprehensive picture of the roles HARs/HGEs play amongst diverse fetal cell types of the developing human brain.

## Background

Humans have complex social and cognitive behaviors, which are at least partly organized during neurodevelopment, resulting in a brain that is exceptional among primates in terms of size, cortical organization, and connectivity^1–4^. One approach to identifying genomic regions that contribute to human neurodevelopment is to compare human genomic sequences and epigenetic features with other species. Human Accelerated Regions (HARs) are genomic regions that are highly conserved among vertebrates but have higher than expected substitution rates on the human specific lineage of the evolutionary tree^5–10^. The fact that these regions are conserved in vertebrates suggests that they are functionally important regions (such as regulatory regions), and the high substitution rate in the human lineage suggests that their function has been modified specifically in humans^4^. Human Gained Enhancers (HGEs) and Human Lost Enhancers (HLEs) are genomic elements that have been identified by examining changes in epigenetic features that mark regulatory activity, such as H3K27 acetylation, between primate and human brains in both fetal and adult contexts^11, 12^.

Massively Parallel Reporter Assays (MPRAs) have shown that many of these HARs function as enhancer regions, with many validated enhancers showing activity during embryonic development^13, 14^. Additionally, HAR specific sequencing in patients with Autism Spectrum Disorder (ASD) coupled with chromatin interaction sequencing (Hi-C) and MPRAs have shown that mutations in HARs that serve as active enhancers may disrupt social and cognitive function^15^. However, understanding the role of these HARs or HGEs across the different human brain cell types present during neuronal development remains a challenge as these genomic regions are mostly in noncoding areas of the genome. Moreover, variation in genomic coding and non-coding sequences between individuals and within individuals (single cell somatic genomic mosaicism) can alter DNA sequences, creating additional challenges in understanding HARs and HGEs^16, 17^. Bulk chromatin accessibility maps of the germinal zone and cortical plate integrated with chromatin interaction maps (Hi-C) have revealed that genes linked to Human Gained Enhancers (HGEs) were enriched for expression in outer radial glia (oRG)^18^. Integrating those same chromatin interaction maps of the human germinal zone and cortical plate with single-cell RNA-seq profiling of human brain development enabled the identification of possible neurodevelopmental risk genes expressed in specific developmental cell types that are regulated by HARs/HGEs^19^. Nevertheless, to our knowledge, there is still no study that directly assays the cell type specific activity of HARs/HGEs in the cells from developing human cortex. Additionally, prior studies focused on the genes regulated by HARs/HGEs but not the transcription factors (TF) binding to these genomic regions that drive human specific gene regulatory networks.

To address these limitations, we aimed to characterize both the upstream transcription factors that bind to HARs/HGEs as well as the downstream genes regulated by these genomic regions in a cell type specific manner by integrating multiple single cell ‘omics datasets. This included data we generated using both a single cell chromatin accessibility assay (scTHS-seq) and single nucleus RNA-seq assay (snDrop-seq) on the same nuclei isolated from the fetal human frontal cortex during weeks 16.6 - 18.2. To further strengthen these analyses, we included recently published fetal human atlases of chromatin accessibility (sciATAC-seq) and gene expression (sciRNA-seq) generated from across the fetal human cortex during weeks 12.7 – 17.8^20, 21^. We also analyzed a previously published scTHS-seq dataset from the human adult visual cortex^22^ to assess for differences between fetal and adult cell types. With these integrated analyses, we were able to look for cell types exhibiting higher than expected accessibility of HARs/HGEs, as well as TFs with binding motif enriched within accessible HARs/HGEs. Integration of fetal brain datasets with chromatin conformation (Hi-C) maps from the developing human cortical cell types further permitted the construction of HAR/HGE-centered TF regulatory networks, with each TF linked to a gene via one or more HAR/HGE^23^.

## Results

### Single cell chromatin accessibility profiling of the developing human cortex

To characterize the chromatin accessibility landscape of the developing human fetal cortex, we used scTHS-seq (scTHS)^22^ to generate profiles from 40,678 single nuclei from 16.6 and 18.2 week fetal frontal cortices **(Figure 1a, Table S1, Methods)**. We additionally profiled 7,865 nuclei from the same samples using snDrop-seq (snDrop), providing gene expression profiles that would permit more confident identification of molecular cell types within the chromatin data^22, 24^ **(Figure 1a, Table S1, Methods)**. We then developed a computational pipeline using these multi-modal datasets to identify cell types with higher-than-expected accessibility of genomic regions linked to human specific evolution **(Figure 1a)**. This includes integration with published Hi-C data and transcription factor motif databases to identify regulatory networks affected by human specific genomic regions^5–11, 23^.

**Figure 1:**
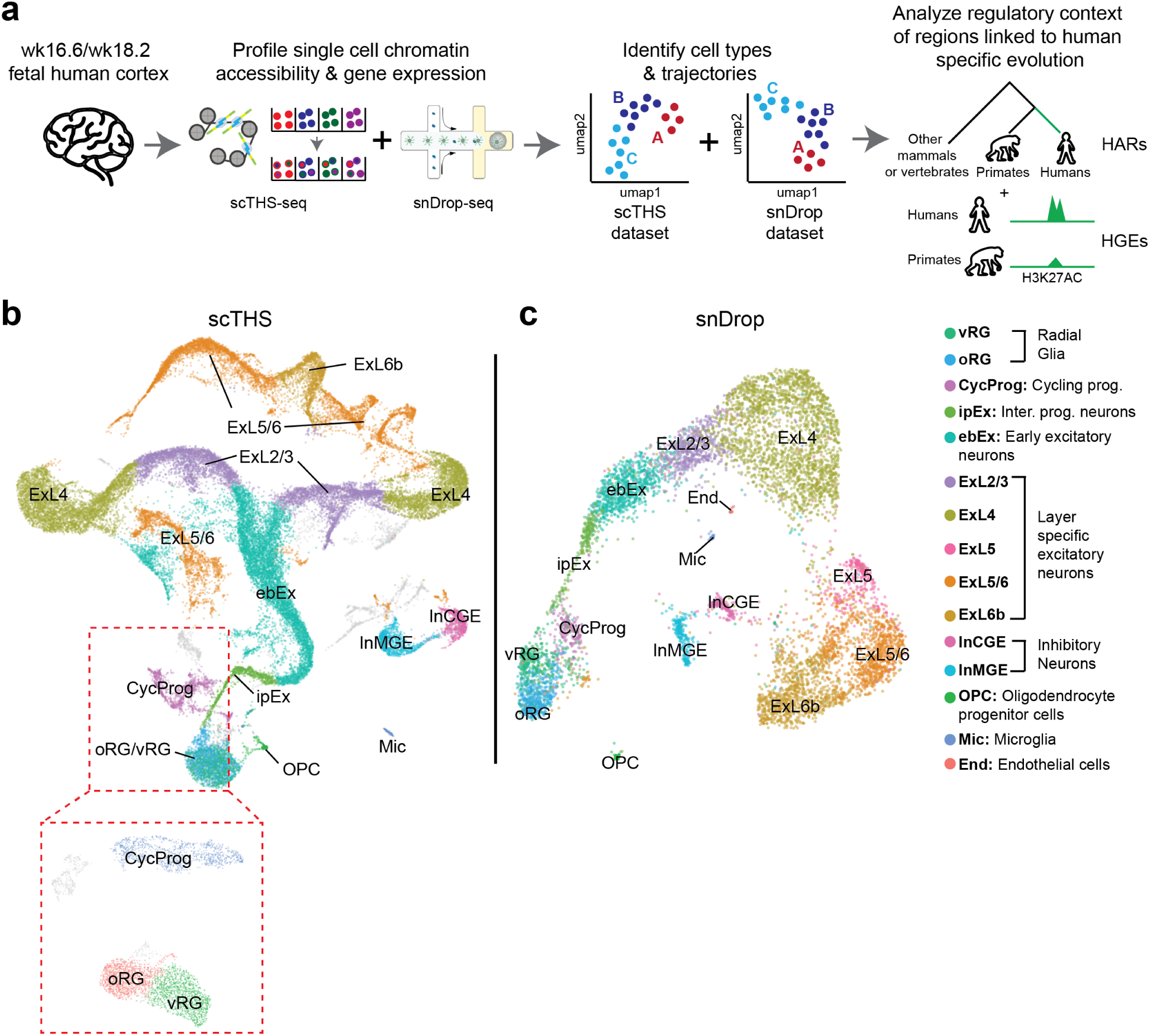
Single cell chromatin accessibility profiles of the developing human cerebral cortex. **a)** We dissociated nuclei from gestational week 16.6 and 18.8 human cortices and used single cell THS-seq (scTHS) to profile chromatin accessibility and single nucleus Drop-Seq (snDrop) to profile gene expression. We identified cell types, and analyzed the cell type specific accessibility patterns of genomic regions linked to human specific evolution. **b)** UMAP embedding of our scTHS chromatin accessibility dataset. **c)** UMAP embedding of our snDrop gene expression dataset.

To analyze our scTHS data, we identified a consensus set of 434,899 accessible regions and generated a matrix of regions by cells^25^ **(Table S1, Methods)**. We then used CisTopic to identify the primary components in the chromatin data, clustered the cells using those components, and generated a predicted gene activity matrix to facilitate cell type identification ^26–28^ **(Methods)**. To analyze our snDrop data, we generated a gene expression matrix, clustered the cells, and identified key marker genes for each cluster^24, 27, 29^ **(Methods)**.

To annotate cell types in our snDrop dataset, we matched cluster specific marker genes to canonical markers from previous scRNA-seq studies of the human fetal cortex and confirmed these annotations through correlation of average gene expression values to reference RNA cell types^30^ (**Figure S1a, Methods)**. We then correlated the average gene expression of our snDrop cell types with the average gene activity of our scTHS clusters to generate a rough map of chromatin cell type identities **(Figure S1b, Methods)**. The activity levels of these same canonical marker genes^30^ **(Figure S1c, Methods)**, as well as the correlation values between RNA and chromatin cell types **(Figure S1d)** further supported our chromatin cell type assignments.

scTHS chromatin and snDrop RNA data both independently generated UMAP embeddings that were consistent with the developmental trajectory of excitatory neurons. This involved the progression from radial glia cell types (**oRG**, **vRG**) through intermediate progenitors (**ipEx**) and early born excitatory neurons (**ebEx**) to the more mature layer specific excitatory neurons (**ExL2/3**, **ExL4**, **ExL5/6**, **ExL6b**) ^31, 32^ **(Figure 1b, 1c)**. Other cell types that were visualized outside of this trajectory were the inhibitory neurons, which migrate into the frontal cortex from the Medial Ganglionic Eminence (**InMGE**) or the Caudal Ganglionic Eminence (**InCGE**), as well as Oligodendrocyte progenitors (**OPC**) and Microglia (**Mic**)^30^ **(Figure 1b, 1c)**. Endothelial cells (**End**) were a rare cell type detected in the snDrop, but not the scTHS, dataset, possibly due to our scTHS chromatin analysis collapsing the Endothelial and Microglia (**Mic**) cell types as chromatin accessibility profiling typically does not separate out cell types as well as gene expression profiling^22^ **(Figure 1b, 1c)**. To visually separate Outer and Ventral Radial Glia (**oRG** and **vRG**), we selected only **oRG**, **vRG**, and **CycProg** cells from our binary chromatin accessibility matrix and reran LDA and UMAP using CisTopic **(Figure 1b, Methods)**.

To assess the reproducibility of our analyses, we additionally examined a published dataset profiling human fetal development that the entire fetal cortex (sciATAC/sciRNA) **(Methods)**^20, 21^. We found a lack of radial glia (**RG**) in the sciATAC/sciRNA cell type annotations, which was unexpected given that the underlying samples were obtained from a period of active neurogenesis (weeks 12.7 – 17.8)^33^. We also found a large excitatory neuron (**Ex**) cluster that was capable of being further sub-clustered to resolve additional key developmental subtypes, such as migrating **ebEx** neurons **(Methods)**^20, 21^. For these reasons, we sought to re-assess the sciATAC/sciRNA cell type annotations based on correlation with the average expression of reference developmental cortex cell types and adult snDrop cell types^22, 30^ **(Figure S2a, S2c, Methods)**. We further visualized key **RG** (*GLI3, VIM, NES, FABP7, SOX2*), **ipEx** (*EOMES, PPP1R17, NEUROG2, PAX6*) and astrocyte (**Ast**) (*S100B, NEU1*) markers **(Figure S2b, S2d)**. Using this approach we were able to re-annotate cell types within this published data set, finding radial glia, intermediate progenitor and excitatory neuron subpopulations that better correspond with expected marker gene profiles **(Figure S2a-d, Methods)**. Robustness of the re-labeled sciATAC and sciRNA cell types was evident from their average marker gene expression or activity profiles **(Figure S2e)**, as well as the expected trajectories observed from the UMAP embeddings **(Figure S2f)**. Therefore, we find a high correspondence in the **RG** to **Ex** trajectories between the sciATAC/sciRNA datasets and our own scTHS/snDrop datasets **(Figure S2f)**. Through this we were able to compile two multi-model data sets with harmonized cell type populations underlying a similar temporal window of human brain development.

### The cell type specific activity of human specific genomic regions in the developing human cortex

Most Human Accelerated Regions (HARs) and Human Gained Enhancers (HGEs) reside in noncoding regions of the genome, with only 5% – 15% of regions overlapping an exon, suggesting these elements are mostly noncoding regulatory regions **(Figure S3a, Table S2)**. Thus, understanding the molecular functions of HARs/HGEs requires profiling the activity of the noncoding genome as well as the regulatory context in which HARs/HGEs operate. To this end, we used the multi-model datasets to identify both the cell types where these HARs/HGEs were most active, and the transcription factors (TFs) with motifs enriched in highly active HARs/HGEs **(Figure 2a)**. We also included adult accessible chromatin from the visual cortex that was profiled using scTHS-seq to assess how fetal cell types differ from those in the adult^22^.

**Figure 2:**
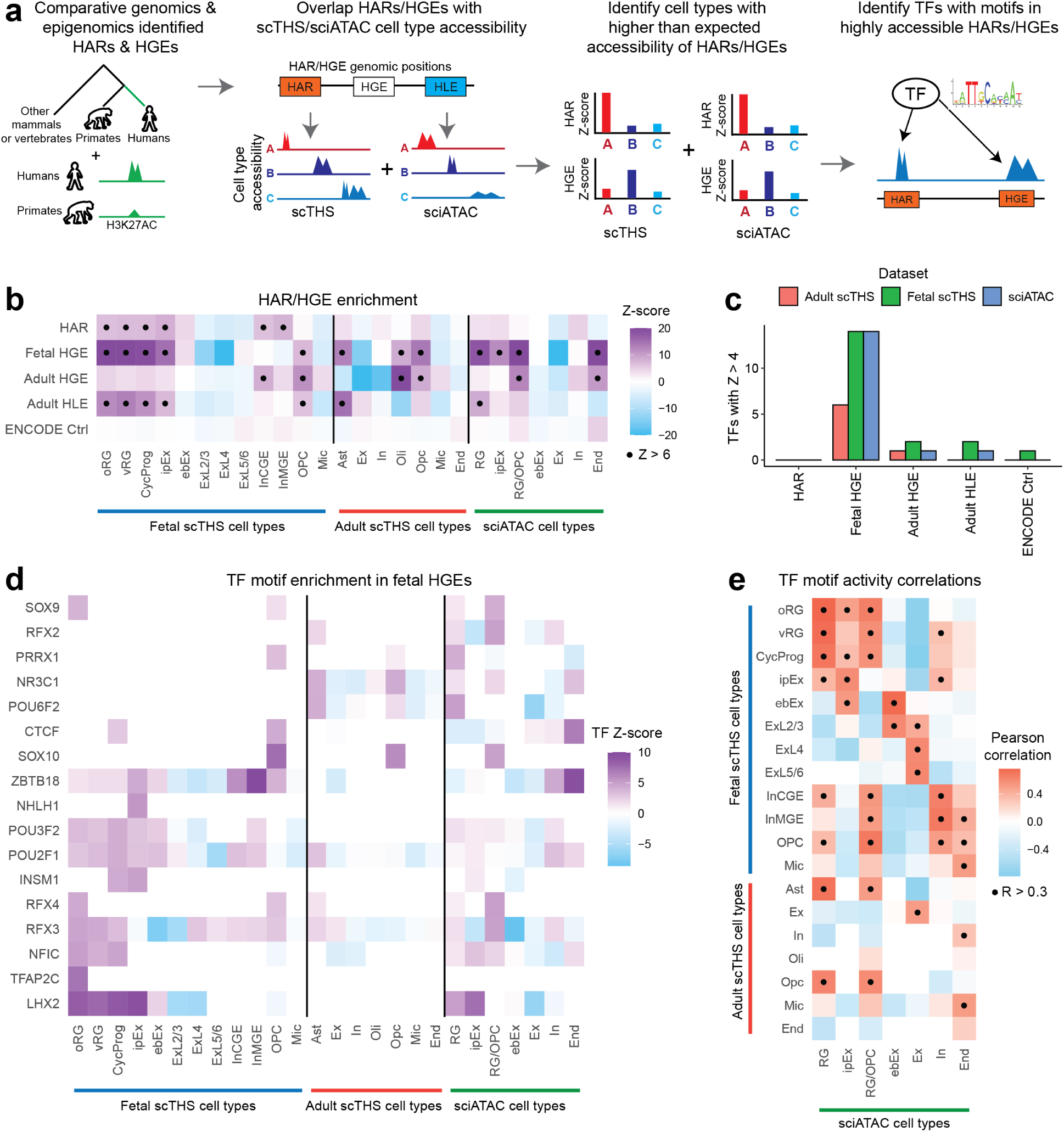
The cell type specific activity of human specific genomic regions in the developing human cortex. **a)** We compiled Human Accelerated Regions (HARs) and Human Gained Enhancers (HGEs) and identified cell types with higher than expected accessibility at those genomic regions. We then found transcription factors (TFs) with motifs enriched in accessible HARs/HGEs. **b)** Heatmap of HAR/HGE cell type enrichment across our fetal/adult scTHS cell types and sciATAC cell types. We sampled 20,000 random DNase I accessible sites as a set of control genomic regions. Dots represent enrichment Z-scores greater than 6. **c)** Barplot of the number of TFs with motif enrichment Z-scores greater than 4 in at least one cell type for each dataset. **d)** Heatmap of cell type specific TF motif enrichment in fetal HGEs. We visualized the top 5 TFs with an enrichment Z-score greater than 4 for each cell type. **e)** Pearson correlations of TF motif enrichment in fetal HGEs between our fetal/adult scTHS cell types and sciATAC cell types. Dots represent Pearson correlations greater than 0.3.

We used chromVAR to identify the cell types with greater than expected levels of HAR/HGE accessibility, using 20,000 randomly sampled DNase I accessible sites across all ENCODE cell lines and tissues as a set of control regions^22, 34, 35^ **(Figure 2b, Methods)**. We found that developmental progenitors, including **RG/oRG/vRG** and **ipEx**, were highly enriched for HAR/HGE accessibility, which was not the case for mature excitatory neurons **(Figure 2b)**. Astrocytes (**Ast**), which have been previously linked to human cortical expansion, and both fetal and adult oligodendrocytes (**Oli/OPC**) were also enriched for HAR/HGE accessibility^36^ **(Figure 2b)**. While fetal HGEs showed the strongest overall enrichment/depletion, most of the HARs/HGEs were enriched in related cell types, consistent with their tendency to converge within similar cells^19^ **(Figure 2b)**. Overall, neuronal progenitors and glial cells were more associated with human specific genomic activity, with fetal HGEs tending to have the highest level of overall activity in these cell types.

With the identification of cell types enriched for global HAR/HGE activity, we next sought to identify specific transcription factors (TFs) that might bind within these genomic regions. This would indicate whether human specific sequence acceleration or human specific epigenetic activity influences the regulatory activity of these TFs through modulating access to their binding sites. We used chromVAR to find transcription factors (TFs) with motifs enriched in HARs/HGEs that showed high levels of cell type specific activity, or with strong enrichments across the different types of HARs/HGEs **(Figure 2c, Table S2, Table S3, Methods)**. Fetal HGEs were the only set of genomic regions with a significant number of TFs passing our enrichment Z-score threshold (Z > 4) **(Figure 2c, Methods)**. These regions gained epigenetic activity in human fetal brain cell types, which could be the result of sequence changes elsewhere or a result of changes in cell type composition in the fetal human cortex versus the primate cortex^11^. Since fetal HGEs also had the strongest overall cell type activity, we focused the remainder of our analyses on these regions **(Methods)**.

For each cell type enriched for fetal HGE activity, we visualized the top TFs that pass our motif enrichment threshold **(Figure 2d, Methods)**. As with the overall fetal HGE enrichment, most of the TF activity occurred in neuronal progenitors and glial cells **(Figure 2d)**. Many of the TFs with strong motif enrichment in fetal HGEs, such as *LHX2*, *RFX3*, and *POU3F2*, have been previously linked to brain development and/or psychiatric disorders^37–40^. *LHX2* is a particularly interesting TF as it shows strong fetal HGE motif enrichment in neuronal progenitors (**RG**, **ipEx**) across both datasets, and is known to be a key regulator of forebrain development^37^. Prior studies have also found that an *LHX2* enhancer showed accelerated evolution in placental mammals and was critical to social stratification in mice^41^. Thus, by looking for cell type specific TF motif enrichment in fetal HGEs, we found that these regions have the potential to influence forebrain development and social behavior by activating *LHX2* binding sites in neuronal progenitors.

To assess the reproducibility of our TF analysis, we correlated the TF motif enrichment Z-scores in fetal HGEs across scTHS and sciATAC cell types **(Figure 2e, Methods)**. From this we found that corresponding cell types, like neuronal progenitors (**RG**/**ipEx**), showed highly correlated motif enrichments across datasets. Interestingly, *LHX2* had the strongest TF motif enrichment in both the sciATAC **RG** and scTHS **oRG** cell types, which further implicates involvement for *LHX2* regulatory activities in fetal HGE-dependent effects on human development **(Figure S3b)**. Prior studies have shown that LHX2 knockouts in mice result in early neurogenesis and that primate specific **oRG** cells have extensive self-proliferative abilities, suggesting that the size of the **RG/oRG** progenitor population at the time of neurogenesis could contribute to cortical size^42, 43^. Overall, by correlating the fetal HGE TF motif enrichments across our cell types and the fetal atlas cell types, we find that the motif enrichment analysis is both reproducible and cell type specific.

As further confirmation of our findings, we examined how our HAR/HGE TF binding site enrichments corresponded to differentially accessible regions (DARs) identified from chimp to human from a recent organoid study^44^ **(Figure S3c, Methods)**. A strong correlation in the scTHS/sciATAC cell type specific HAR/HGE TF motif enrichments with TF motif enrichment in the human/chimp DARs **(Figure S3c)** suggest that human/chimp DARs converge with HARs/HGEs in terms of their TF motif activity patterns. Further correlations of fetal HGE from **vRG** and **Ast** with human/chimp DARs again identified LHX2 motifs as the primary driver of the RG TF motif correlation **(Figure S3d).** Alternatively, the **Ast** correlation was found to be driven by several motifs including those for POU6F2 and FOXG1 **(Figure S3d)**. Therefore, we demonstrate a correspondence between TF binding motifs identified from our analyses with those enriched in human over chimp accessible regions identified from an orthogonal dataset. This provides additional support for an expanded LHX2 regulatory role in human neural progenitors.

### Capturing the transcription factor regulatory context of fetal HGEs

After characterizing the cell type activity of HARs/HGEs and identifying enriched TF motifs, we next sought to build HAR/HGE-centered regulatory networks. These networks link upstream TFs that bind to HARs/HGEs with downstream genes whose promoters are in contact with HARs/HGEs **(Figure 3a)**. Our goal with these networks was to provide a model of human evolution where these human specific genomic regions modify TF regulatory networks by modulating connections between TFs and the genes they regulate. We again focused on fetal HGEs, and first linked TFs to fetal HGEs using their binding motifs **(Figure 3a, Table S4)**. However, the presence of a TF motif in a HGE does not necessarily mean that TF is bound to that region in a given cell type. Thus, for each TF, we only linked a TF to HGEs if the overall TF motif activity was also correlated with HGE accessibility, and where TF expression was also detected **(Figure 3a, Methods)**. We then linked fetal HGEs to downstream genes using published chromatin conformation capture (Hi-C) data generated from **RG**, **ipEx**, **ebEx** and **In** cell populations, keeping only those HGEs that were also accessible in these cell types **(Figure 3a, Table S4, Methods)**^23^. By linking TFs to HGEs, and HGEs to genes, we built HAR/HGE-centered regulatory networks where TFs regulate genes via binding to fetal HGEs **(Figure 3a)**. To assess reproducibility in these networks, we built them independently from both the scTHS and sciATAC **(Figure 3a).** Therefore, we generated two independent, yet corroborating HAR/HGE networks through co-assessment of the accessibility, expression, and correlation of TFs and HGEs in the developing fetal brain cell types **(Table S4)**.

**Figure 3:**
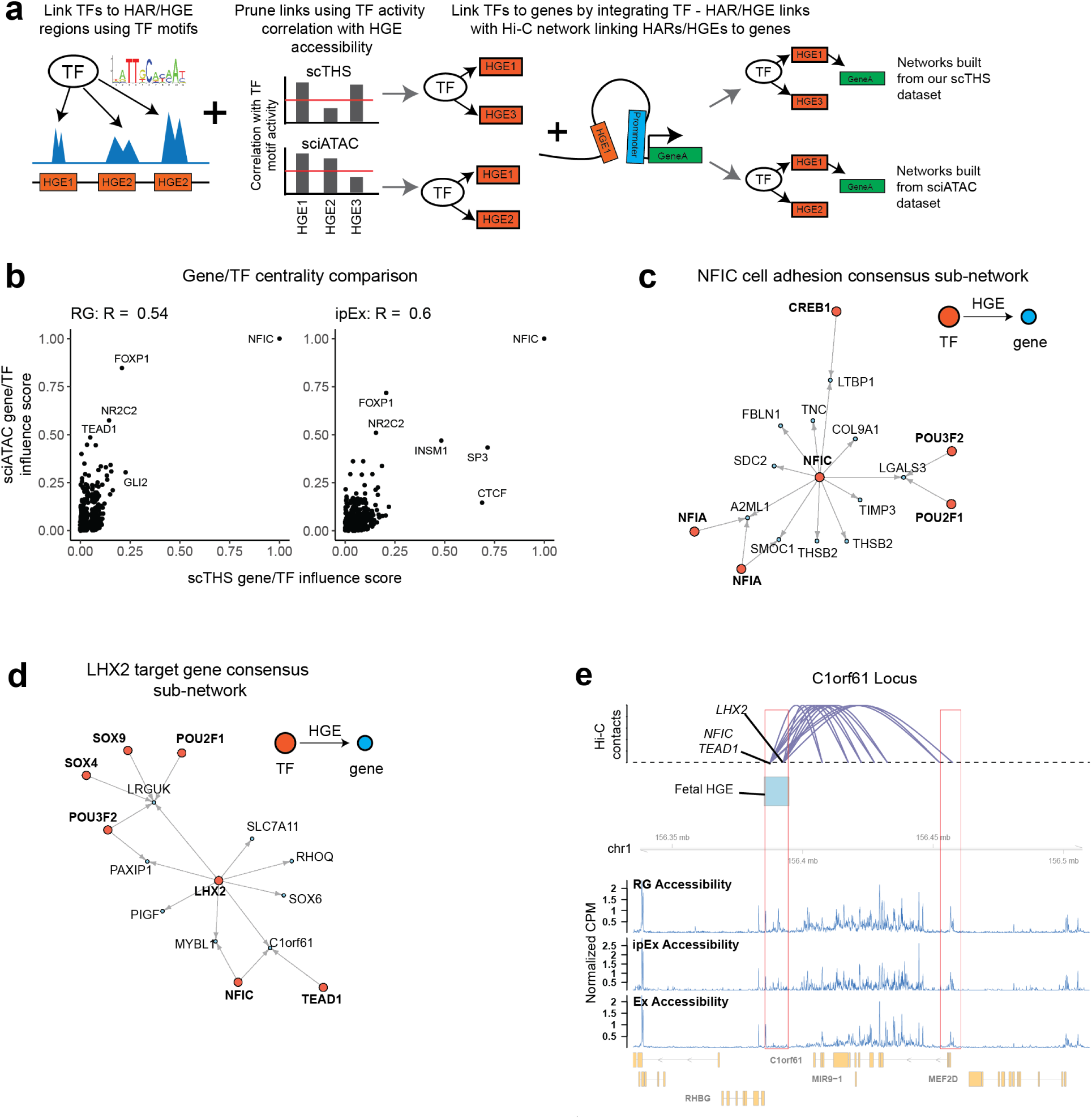
Capturing the transcription factor regulatory context of fetal human gained enhancers. **a)** We linked TFs to HGEs if the TF motif was in the HGE, pruning motif – HGE links by correlating single-cell TF motif activity with HGE accessibility using our fetal (scTHS) dataset and the fetal atlas (sciATAC) dataset. We then used Hi-C networks to link accessible HGEs to gene promoters, and built TF regulatory networks where TFs regulate genes via binding to HGEs. We generated networks using both our scTHS dataset and the fetal atlas sciATAC dataset. **b)** Scatterplots visualizing the influence of nodes (genes & TFs) in each network by plotting eigenvector centralities across the sciATAC and scTHS networks. Eigenvector centrality finds nodes that point to other nodes who are themselves highly connected, thus allowing for a more accurate measurement of node influence than simply counting the number of connections for each node. **c)** Force directed layout of a neural progenitor (**RG/ipEx**) *NFIC* cell adhesion network consisting of all *NFIC* target genes related to ECM and cell adhesion present in both sciATAC and scTHS networks. **d)** Force directed layout of a neural progenitor *LHX2* network consisting of all *LHX2* target genes present in both sciATAC and scTHS networks. **e)** Visualization of the fetal HGE region that contains *LHX2* and *NFIC*/*TEAD1* motifs and is in contact with the C1orf61 promoter. Arcs represent a chromatin contact from the neural progenitor Hi-C networks. Tracks showing the normalized accessibility of the locus in Counts per Million (CPM) are displayed for **RG**, **ipEx** and mature **Ex** cell types.

To understand the size and connectivity of these networks, we plotted the number of nodes and edges for each network as well as the number of overlapping nodes and edges between corresponding scTHS and sciATAC networks **(Figure S4a)**. The sciATAC networks tended to have more nodes (TFs/genes) and edges (TF – gene links) than scTHS networks, possibly due to the higher sampling depth for the sciATAC assays **(Figure S4a)**. As the sciATAC networks tended to be larger than the scTHS networks, we plotted the fraction of scTHS network nodes and edges also found in the sciATAC networks **(Figure S4b)**. Roughly 60% of scTHS network nodes and 20% of scTHS network edges were also in sciATAC networks, and neuronal progenitor networks (**RG** & **ipEx**) tended to have a greater overlap of nodes/edges than **Ex** or **In** networks **(Figure S4b)**. Since **Ex** and **In** cells were not enriched for overall fetal HGE accessibility and their corresponding networks tended to be fairly sparse, we focused the remainder of our analysis on the human neuronal progenitor (**RG** & **ipEx**) networks.

We next sought to identify influential TFs and genes in each network. To this end, we computed the eigenvector centrality for nodes (TFs or genes) in each network, which assigns an influence score to each node (a TF or gene) based on how many other highly connected nodes it is linked to **(Methods)**. Measuring eigenvector centrality enables us to find influential nodes that may not themselves be highly connected, but are linked to other highly connected nodes **(Methods)**. We then correlated the centralities across corresponding sciATAC and scTHS networks^45^ **(Figure 3b)**. Overall, the centralities for corresponding scTHS and sciATAC neuronal progenitor networks were strongly correlated, and the outsize influence of Nuclear Factor I/C, or NFIC, in both **RG** and **ipEx** was found to be the primary driver for these correlations **(Figure 3b)**. NFIC’s influence on these progenitor networks was especially interesting as the Nuclear Factor I (NFI) family of TFs are known to play key roles in development^46^.

Given the influential role for NFIC in neuronal progenitor networks, we assessed NFIC target genes in greater depth. Our motif binding analysis found that NFIC was bound to 1,624 fetal HGEs across all scTHS networks and 1,976 fetal HGEs across all sciATAC networks **(Methods)**. We found a significant overlap in NFIC gene targets across the scTHS networks and the corresponding sciATAC networks **(Figure S4c)**. To find pathways enriched in NFIC target genes, we compared the genes against a combination of the Hallmark pathways and the Canonical pathways from the Molecular Signatures Database^47, 48^ **(Figure S4d)**. Pathways that showed a strong enrichment across both sciATAC and scTHS networks were associated with Notch Signaling and Matrisome/ECM Glycoproteins **(Figure S4d)**. Notch signaling is critical to numerous aspects of brain development, including maintenance of neural stem cells^49^. Furthermore, the NFI family was found to regulate Notch signaling in Glioblastoma and **Ast** development, suggesting that NFIC may in part regulate human cortical development via Notch signaling^50, 51^. The enrichment for Matrisome/ECM pathways also suggests that NFIC may be affecting cell adhesion, a critical cell property that affects neuronal migration and development^52^. The NFI family also regulates cell adhesion molecules during granule cell development, affecting cell migration, axon outgrowth, and dendrite formation^53^. Overall, pathway enrichment analysis identified Notch signaling and cell adhesion pathways, both expected to play key roles in neuronal differentiation and maturation, as modulated by NFIC via active human gained enhancers.

To further investigate how fetal HGEs modulate NFIC’s regulation of cell adhesion, we built a consensus sub-network of all NFIC target genes in the Matrisome and ECM gene sets that are present in both sciATAC and scTHS networks **(Figure 3c, Methods)**. Both NFIC and NFIA target *SMOC1*, encoding a calcium binding protein commonly found in basement membranes^54^ **(Figure 3c)**. Mutations in *SMOC1* cause developmental abnormalities, including forebrain defects^54^. In mice, Nfic co-localizes with Pbx to regulate cortical patterning and *Smoc1* expression in the ventricular zone, suggesting that modifications to the relationship between NFIC and *SMOC1* via HGEs could be relevant to human specific brain development^55^. NFIC, POU3F2, and POU2F1 all target the *LGALS3* gene, the protein product of which can mediate cell adhesion via interactions with glycosylated proteins, has been implicated in the proper development of locomotive function, and has been linked to Huntington’s disease^56, 57^. Thus, our NFIC target gene network identified SMOC1 and LGALS3 as potential modulators of cell adhesion whose expression may be influenced by HGEs that activate NFIC binding sites.

Since LHX2, a transcriptional activator, had consistently strong motif enrichment in fetal HGEs and human/chimp DARs, we also visualized the LHX2 target gene sub-network **(Figure 3d)**. The sub-network showed that LHX2, NFIC, and TEAD1 all regulate expression of the *C1orf61/CROC-4* gene, encoding a brain specific cDNA that is involved in cellular remodeling of brain architecture via the c-fos signaling pathway^58^ **(Figure 3d)**. We also found LHX2 regulates the *SOX6* gene, a relationship that had been previously documented in retinal development^59^ **(Figure 3e).** Upon closer examination of the *C1orf61/CROC-4* locus, we identified a high number of chromatin interactions between a single HGE site and several other sites within the *C1orf61/CROC-4* gene body, suggestive of alternative *C1orf61* promoters **(Figure 3e)**. We also found significant motif co-occupancy between LHX2 and NFIC, which was also found during retinal development^60^ and indicative of possible cooperation between these TFs. Overall, our findings implicate LHX2 and NFIC, both of which have shown strong motif enrichment and network influence in our analyses, as being highly positioned for key roles in modulating human specific brain development.

We also looked for influential connections and genes in neuronal progenitor networks using edge betweenness centrality, which looks for connections between nodes that act as bridges between different densely connected sub-networks^61, 62^ **(Figure S4e, Methods)**. The connection between NFIC and GLI2 was by far the most influential in both RG networks, driving the correlation between the sciATAC and scTHS **RG** networks. GLI2 is a key TF in the Hedgehog signaling pathway that regulates patterning during brain development^63^ and while NFIC has been previously linked to the Hedgehog signaling cascade in the context of tooth root development, our edge betweenness analysis suggests this NFIC – Hedgehog signaling cascade may also be related to human specific brain development^64^. Unlike in **RG** networks, the correlation between sciATAC and scTHS in **ipEx** networks was driven by a number of edges, including an edge linking NFIC and ZNF423, a key regulator of cell cycle progression in neuronal progenitors^65^ **(Figure S4e)**. The influence of fetal HGEs on NFIC’s regulation of cell cycle progression is consistent with the cycling properties of **ipEx** cells^43^.

Since the NFIC – GLI2 edge had disproportionate influence on the RG networks, we examined the locus containing the relevant fetal HGEs and the *GLI2* promoter **(Figure S4f)**. Interestingly, this locus had a high density of fetal HGEs overall, which further suggests a role for GLI2 in human specific brain development **(Figure S4f)**. The two fetal HGEs contacting the *GLI2* promoter contain NHLH1/GLI2 motifs and NFIC/FOXP1/SREBF2 motifs, respectively suggesting an autoregulatory loop **(Figure S4f)**. The NFIC/SREBF2 motif co-occupancy has been shown to collaborate in regulating a cholesterol synthesis pathway, suggesting that this co-occupancy may play a role in human specific development^66^ **(Figure S4f)**. Both the *GLI2* promoter and the two fetal HGEs contacting the promoter have higher accessibility in neuronal progenitors (**RG** & **ipEx**) than mature **Ex** cells, confirming that these regions are primarily active in neuronal progenitors **(Figure S4f)**. Therefore, we found that enrichment of differentially accessible fetal HGEs within the GLI2 promoter that may drive additional human-specific expression of this factor within progenitor cells. Careful examination of these fetal HGEs further finds both a potential auto-regulatory function of GLI2, as well as a possible cooperative role for NFIC and SREBF2.

## Discussion

We used single-cell chromatin accessibility and gene expression profiles from the developing fetal human cortex to study both the upstream TFs binding to HARs and HGEs, as well as their candidate downstream regulated genes. Accessibility within these genomic regions was highest in neuronal progenitors and glial cells, indicating that HAR/HGE regulatory activities primarily target progenitor populations. Consistent with this high activity in progenitors, we found that fetal HGEs had the strongest overall cell type enrichment. We also found that LHX2, a key regulator of forebrain development, had significant enrichment of its binding motifs within highly accessible fetal HGEs specifically in neuronal progenitors. Then, by using Hi-C networks to link TFs to downstream genes via fetal HGEs, we created HGE-centered regulatory networks. This analysis identified a central regulatory role for NFIC within these neuronal progenitor regulatory networks, and found that NFIC target genes were enriched for Notch signaling and cell adhesion pathways. A key NFIC target gene in the cell adhesion pathway was *SMOC1*,which encodes a basement membrane protein key to numerous aspects of embryonic development^54^. These networks also identified potential TF cooperation, such as the co-occupancy of LHX2 and NFIC motifs. Thus, we were able to create a more holistic view of the biological function of HARs/HGEs by analyzing both the TFs binding to these regions and the genes regulated by those TFs via HARs/HGEs in a cell type specific context.

By using single cell chromatin accessibility, we were able to directly measure HAR/HGE accessibility in specific cell types, whereas previous work either measured bulk HAR/HGE accessibility or linked HARs/HGEs to genes for cell type specific interpretation^18, 19^. Matched single cell gene expression datasets enabled us to filter for TFs that were expressed in the same cell types showing HAR/HGE motif enrichment. This multi-omics approach also allowed for fewer false positive contacts in the Hi-C data between HARs/HGEs and downstream genes, as we were able to filter for HARs/HGEs and genes that are accessible or expressed in the same cell types. Thus, the integrated use of chromatin accessibility, gene expression, and Hi-C modalities enabled us to construct high quality regulatory networks.

However, when building the TF regulatory networks, we relied on the presence of TF motifs in HARs/HGEs to link those genomic regions to TFs. This can lead to false positive TF – HAR/HGE links since the presence of a TF motif does not necessarily mean the TF is bound to that genomic region^67^. To reduce these false positive links, we correlated each TF’s motif activity with the accessibility of HARs/HGEs that contained the TF’s motif; this ensured that we only identified HARs/HGEs that were accessible when the TF motif tends to be active. A benchmarking study of various features used to predict TF binding has shown that chromatin accessibility and binding motif presence have the best predictive value relative to ground truth Chip-Seq datasets^68^. Thus, ensuring that the HARs/HGEs with TF motifs are accessible in the same cells that the TF is active should improve our ability to link TFs with HARs/HGEs.

While there are existing computational tools for predicting transcription factor binding, they have almost exclusively been trained on Chip-Seq datasets from cell lines and may not generalize to our fetal brain data^69^. Future work applying Chip-Seq or Cut & Run to human and chimp organoids or teratomas at different stages of organoid development or FACS sorted progenitor cell types could enable us to directly measure transcription factor binding during human and/or primate development^70, 71^. However, these methods require a unique antibody for each TF, making it difficult to assay TFs in a high throughput manner. A higher throughput strategy could be to run Chip-Seq on organoids/teratomas or sorted human progenitor cell types using a set of training transcription factors, and then using these transcription factor binding site prediction methods to build a comprehensive database of predicted transcription factor binding in these models of the fetal human brain.

To ensure biological reproducibility of our HAR/HGE enrichment and TF motif analysis results, we used independent single cell chromatin accessibility datasets generated from two different methods. However, the TF regulatory networks built from these two datasets were not entirely independent as they used the same Hi-C dataset as well as the same motif position database. Nevertheless, it was helpful to analyze these networks separately given dataset specific elements present within each TF regulatory network, such as the accessibility of HGEs in any given cell type or the correlations between TF motif activity and HGE accessibility. We also compared our results with a published study of single cell gene expression and accessibility in human and chimp organoids, confirming that our HAR/HGE analysis correlated well with human/chimp differential accessibility. Thus, by using two chromatin accessibility datasets as well as an orthogonal dataset comparing human and chimp organoids, we were able to ensure reproducibility of our analyses.

Region specific differences and differences in gestational time points could also explain some of the discrepancies between the sciATAC and scTHS TF networks. Our scTHS data was sampled from the fetal frontal cortex of two samples from weeks 16.6 and 18.2, while the sciATAC data was sampled from the entire fetal cortex of 121 samples from weeks 12.7 to 17.8. Additionally, our adult scTHS dataset was specifically from the visual cortex. Variation in the genome between individuals and within individuals may also explain some of the discrepancies between sciATAC and scTHS datasets^16, 17^. Specifically, single cell somatic genomic mosaicism in the human brain may contribute to changes in HAR/HGE activity between cells and across individuals. Future studies that simultaneously assess genomic sequences, chromatin accessibility, and resulting transcriptomic changes in single cells would clarify these relationships.

Overall, we relied on a Hi-C dataset to link HARs/HGEs to the genes they regulate. However, recent work using CRISPR-QTL frameworks has shown that most enhancer – gene pairs do not show up as Hi-C contact pairs^72^. Thus we are most likely missing the vast majority of genes linked to HARs/HGEs. A future approach to comprehensively identifying the genes that HARs/HGEs regulate could employ a similar CRISPR-QTL framework applied in organoid or teratoma models to more robustly identify the genes regulated by HARs/HGEs^73, 74^.

Additional future work could also look into how the epigenetic changes underlying Human Gained Enhancers (HGEs) arose in the human cortex. One possibility is that mutations in other areas of the genome that regulate epigenetic activity could lead to these HGEs gaining enhancer functionality. For example, a subset of HARs could be responsible for driving the formation of some HGEs and exploring this potential link between HARs and HGEs could lead to a deeper insight into human specific brain development. An alternative possibility is that active enhancer regions could have been copied and pasted into new areas of the genome via transposable elements. Future work could investigate this possibility by looking for transposon sequences near HGEs and then running gain of function experiments in primate organoids.

## Conclusions

Thus, by leveraging a single cell multi-omics approach to assess the biological context of human specific noncoding regulatory regions, we were able to identify candidate TF regulatory networks that may play a role in the evolutionary development of the human brain. These networks directly model the hypothesis that these human specific noncoding regulatory regions modify human specific brain development by modulating the regulatory relationships between TFs and genes. We believe this type of holistic approach to analyzing noncoding regions can help unravel the biology behind not just human specific regulatory regions, but noncoding regulatory regions in general.

## Declarations

### Ethics approval and consent to participate

All human tissue protocols were approved by the Office for Human Research Protection at Sanford Burnham Prebys Medical Discovery Institute and conformed to National Institutes of Health guidelines.

### Consent for publication

Not applicable

### Availability of data and materials

Analysis scripts and functions for running the HAR/HGE enrichment and building HAR/HGE networks can be found at www.github.com/yanwu2014/chromfunks. Scripts for processing raw scTHS-seq data can be found at https://github.com/yanwu2014/scTHS-Seq-processing and scripts for processing raw snDrop-seq data can be found at https://github.com/chensong611/snDrop_prep/. Gene expression and chromatin accessibility matrices and Seurat objects as can be found on our FTP server at ftp://genome-miner.ucsd.edu/fetal_brain_data/. The original sciATAC-seq and sciRNA-seq data from the developing human brain can be found at: https://descartes.brotmanbaty.org/.

### Author contributions

K.Z. and Y.W. developed the conceptual ideas. Y.Y. dissociated the fetal brain tissue into single cells. S.C. ran the snDrop-seq protocol. B.S. and E.D. ran the scTHS-seq protocol. B.L. processed the snDrop-seq data. Y.W., W.D., and S.L. processed the scTHS-seq data. Y.W. reprocessed the sciATAC-seq and sciRNA-seq datasets and ran the transcription factor motif and network analysis. Y.W, B.L., K.Z., J.C., Y.Y., B.S, S.C., and W.D. wrote the manuscript.

## Acknowledgements

We would like to acknowledge Dr. James Knowles from the USC Center of Genomic Psychiatry and Dr. Robert Chow from the USC Zilkha Neurogenetic Institute for providing the fetal human brain tissue samples. We also would like to acknowledge Dr. Prashant Mali from the UCSD Bioengineering Department, the Integrative Genomics Lab members at UCSD for helpful insights and discussions, the Sanford Consortium Flow Cytometry Core for help with sorting nuclei, and the IGM Genomics Center for sequencing these samples.

## Methods

### snDropSeq nuclei preparation

Sections of flash frozen fetal human brain tissue from gestational week 16.6 and 18.2 were obtained from the Robert Chow Lab at USC. These sections included areas from the frontal cortex and ganglionic eminences. Nuclei were prepared with nuclear extraction buffer (NEB) as described previously^22^. Briefly, flash-frozen brain tissue that was sectioned at 50 µm with a cryostat was chopped and mashed with a scalpel and placed in 1 ml of ice-cold NEB with 1 uL DAPI. Nuclei were extracted with 10–12 up-and-down strokes of a glass Dounce homogenizer with a Teflon pestle in 1 ml of NEB. Samples were passed through a 50-µm CellTrics filter into a 15 mL conical tube and then incubated on ice for 10 min. The total volume of each sample was brought up to 10 mL with PBS + 2 mM EGTA and samples were spun for 10 min at 900g, washed in PBS + 2 mM EGTA, and resuspended in 1 mL PBS + 2 mM EGTA supplemented with 1% fatty-acid-free BSA (Gemini). Approximately 250,000 DAPI+ single nuclei were purified by flow cytometry with a FACSAria Fusion (Becton Dickinson) sorter into PBS + 2 mM EGTA supplemented with 1% fatty-acid-free BSA (Gemini), concentrated at 900g for 10 min, resuspended in PBS + 2 mM EGTA supplemented with 0.01% BSA, and then used directly for droplet encapsulation

### snDropSeq Library Preparation and Sequencing

Drop-seq with modifications optimized for nuclei processing was performed as described previously^22^. Before droplet generation, connecting tubing and syringes were coated with 1% BSA to prevent nonspecific binding of nuclei to the surface, and then rinsed with PBS. To reduce nuclei settling, Ficoll PM-400 was added to the nuclei suspension buffer, rather than the lysis buffer. Nuclei were loaded at a concentration of 100 nuclei/µl and coencapsulated in droplets with barcoded beads purchased from ChemGenes Corporation (cat. no. Macosko201110). When encapsulation was complete, the contents of the droplet-collecting Falcon tubes were overlaid with a layer of mineral oil and then transferred to a 72 °C water bath for 5 minutes to lyse the nuclear membranes. We then proceeded to reverse-transcription (RT) and PCR amplification of cDNA as previously described. A total of 12 snDropSeq libraries were prepared, and cDNA from each replicate was tagmented by Nextera XT and indexed with different Nextera index 1 primers. cDNA libraries were pooled and sequenced on an Illumina HiSeq 2500 with Read1CustSeqB^24^ for priming of read 1 (read 1 was 30 bp; read 2 (paired end) was 120 bp).

### snDropSeq Data Processing and Clustering

Paired-end sequencing reads were processed largely as described (http://mccarrolllab.com/wp-content/uploads/2016/03/Drop-seqAlignmentCookbookv1.2Jan2016.pdf), with additional correction steps as previously described^22^. Briefly, reads were filtered to ensure the presence of a polyT and to remove reads with low sequencing quality bases. The right mate of each read pair was trimmed to remove any portion of the SMART adapter sequence or polyT tails. The trimmed reads were then aligned to the human genome (GENCODE GRCH38) with STAR v2.5 with the default parameter settings. Reads that mapped to intronic or exonic regions of genes as per the GENCODE gene annotation were recorded. We applied one further correction step to fix barcode synthesis errors by inserting N at the last base of the cell barcode for reads in which the first 11 bases of the cell barcode were identical and the last T base of UMI was the same. The digital expression matrix was then generated with genes as rows and cells as columns. We assigned UMI counts for each gene of each cell by collapsing UMI reads that had only 1 edit distance.

UMI matrix cell barcodes were tagged by their associated sequencing library batch ID and combined across independent experiments. Mitochondrial genes not expressed in nuclei were excluded, and only UMI counts associated with protein-coding genes were used for clustering analyses. Nuclei with fewer than 300 molecules or more than 5,000 molecules (outliers) were omitted. We normalized molecular counts by using the total number of UMIs as the estimated library size for each cell. Variance normalization and clustering were done with the PAGODA2 package (https://github.com/hms-dbmi/pagoda2) as described previously^29^. Briefly, we selected 2000 overdispersed genes and computed the top 50 principal components (PCs). We generated clusters using PAGODA2 and then imported the gene expression matrix and the PCs into Seurat for UMAP visualization and marker gene visualization. We identified cell types by correlating the average gene expression of our clusters with the average gene expression of cell type from a published fetal human cortex dataset and by visualizing the expression patterns of previously described marker genes for fetal human cortical cell types^30^ **(Figure S1a, S1b)**.

### scTHS-seq Nuclei Isolation

We prepared nuclei for the single-cell THS-seq chromatin accessibility assay using the protocol described in the snDropSeq nuclei isolation section^22^. Briefly, after flow cytometry, nuclei were kept on ice and spun down at 500g for 5 min at 4 °C, after which supernatant was removed and the pellet was resuspended in lysis buffer. Then nuclei were spun down at 500g for 5 min at 4 °C, supernatant was removed, and the pellet was resuspended in tagmentation buffer. At that point the nuclei sample was ready for nuclei counting. A nuclei concentration of ∼2.4 million nuclei/mL was obtained for each sample.

### scTHS-seq Transposome Generation

We prepared transposons for scTHS-seq as previously described^22^. Briefly each transposon consisted of two oligos: the 74-bp barcoded transposon and the 19-bp universal 5′ phosphorylated mosaic end. In total, there were 384 barcoded r5 transposons, each with a unique 6-bp barcode. For the generation of annealed transposons, 10 µL of each 100 µM oligo was added to each well of a 384-well plate (final concentration: 50 µM), incubated at 95 °C for 2 min, cooled to 14 °C at 0.1 °C/s, diluted to 8.4 µM in TE buffer with a final concentration of 50% glycerol, and then stored at −20 °C.

Tn5059 was generated and normalized for activity at Illumina. Transposome complexes were generated freshly for each scTHS-seq run and used within a few days. First, Tn5059 was diluted to 4.2 µM in standard storage buffer (Illumina), and 1 µL was added to each well of 384-well plate. Next, 1 µL of 8.4 µM annealed barcoded r5 transposon was added to each well, and the 384-well plate was incubated at room temperature for 30 min. For custom nXTv2_i7 Tn5059 transposome generation, the annealed nXTv2_i7 transposon (50 µM) was generated and we incubated 7 µM Tn5059 with 10 µM annealed transposon for 30 min at room temperature and then diluted to 0.7 µM Tn5059 transposome complex with standard storage buffer (Illumina).

### scTHS-Seq Tagmentation, Barcoding and Library Preparation

We ran the scTHS-seq assay as previously described. Briefly, we added 4 µL of cell sample to each well of the 384 well-plate with the loaded transposome complex for a total of ∼960 nuclei per well and a final concentration of 0.7 µM Tn5059 r5 transposome complex. To stop the reaction, we added 4.0 µL of 50 mM EDTA to each well and mixed gently five times with the electronic pipettor, and then incubated the mixture at 37 °C for 15 min. We added one volume of cold 2X FACS buffer (2 mM EDTA, 1% BSA in PBS) to each well, and samples were mixed gently three times with the electronic pipettor and pooled into one tube on ice. We spun down the tube, and resuspended the cells in 1X FACS buffer. Next, 75 µL of propidium iodide (PI; eBioscience) was added, and nuclei were sorted by flow cytometry into 96-well plates containing 10 µL of PBS per well at 100 nuclei per well and kept on ice. Doublets were removed on the basis of forward and side scatter plots, and PI-staining events were selected.

Each 96-well plate of nuclei was then processed individually as previously described. Briefly, 11 µL of guanidine hydrochloride was added to each well and mixed by light vortexing. Reactions were purified with AMPure SPRI beads. 10 µL of 1× NEB Taq polymerase was added to each reaction, and the plate was lightly vortexed to resuspend the beads (SPRI beads left in the reaction), after which the reactions were run at 72 °C for 3 min for end fill-in. For in vitro transcription (IVT) amplification, we used the NEB HiScribe T7 high-yield synthesis kit, with incubation at 37°C for 16 hours. For reverse transcription, we added 2.5 µL of 20 µM random hexamers to each reaction, and used the Clontech SMART MMLV reverse transcriptase kit.

Reactions were incubated at 22 °C for 10 min, then 42 °C for 60 min, and terminated at 70 °C for 10 min. To degrade RNA in cDNA–RNA hybrids, we added 1 µL of 0.5 units Enzymatics RNase H to each reaction, vortexed the plate lightly, and incubated the plate at 37 °C for 20 min. For second-strand synthesis, we added first 2.5 µL of 20 µM sss_scnXTv2 to each reaction and lightly vortexed it, then incubated it for 2 min at 65 °C and immediately cooled it on ice.

Then we added 5.9 µL of NEB taq5X to each reaction and incubated it at 72 °C for 8 min. Double-stranded cDNA fragments then underwent simultaneous fragmentation and 3′ adaptor addition with a custom nXTv2_i7 Tn5059 transposome. To 7-µL volumes of each sample, we added 2 µL of 5× tagmentation buffer, followed by 2 µL of prepared 0.7 µM custom nXTv2_i7 Tn5059 transposomes (final transposome concentration of 0.14 µM). We added 19 µL of 6.32 M guanidine hydrochloride, for a final guanidine hydrochloride concentration of 4 M, to each reaction and briefly vortexed the sample. We eluted sample off SPRI beads held by the magnetic plate and transferred it to a qPCR plate. Standard Illumina Nextera XT v2 barcoding in an 8 × 12 (i5 × i7) format was performed with qPCR, using custom scTHS-seq i5 indexes and standard Illumina i7 indexes.

For pooling, 2 µL (4 µL or 6 µL if yields were low) of each uniquely barcoded qPCR reaction was combined and size-selection was performed as described^22^. Resultant size-selected libraries were quantified with Qubit and sequenced on an Illumina MiSeq system (50 + 32 + 32 single-end reads) for validation, then on the high-throughput Illumina HiSeq 2500 (50 + 8 + 32 single-end reads) for data generation.

### scTHS-Seq Data Processing

We generated Fastq files for Read1, Index1, and Index2 and identified the reads that map to each unique barcode combination using deindexer (https://github.com/ws6/deindexer) with a zero mismatch stringency. This resulted in a single fastq file per cell barcode combination. After deindexing, we appended the cell barcode combination for each read to the read name, and then re-merged all fastq files for alignment. We aligned the merged fastq file to an hg38 reference genome (GCA_000001405.15_GRCh38_no_alt_plus_hs38d1_analysis_set) using BWA.

We then used the snaptools snap-pre command (with –keep-single = TRUE) to generate a snap file, and the snap-add-bmat command to generate 5 kb bins across the entire genome and create a bins by cells matrix^25^. We then used the SnapATAC processing pipeline to filter out cells with fewer than 3000 unique reads, run dimensionality reduction and clustering on the bins by cells matrix, and used MACS2 to call peaks on the reads from each cluster separately. We merged the peaks from all clusters to generate a consensus list of peaks, and then generated a binary peaks by cells matrix^25^.

### scTHS-seq Dimensionality Reduction, Clustering, and Cell Type Identification

We first filtered the peaks by cells matrix for cells with at least 500 accessible sites to remove potential empty barcode combinations, and less than 20,000 accessible sites to remove potential multiplets. After filtering cells, we used cisTopic to run Latent Dirichlet Allocation (LDA) on the peaks matrix with 30 topics, with the optimal number of topics selected using cisTopic’s model selection functionality^26^. We ran UMAP on the LDA topics to generate a 2D visualization of the data and then imported the LDA topics into Seurat v3 and clustered the cells using default parameters^27^. To help with cell type annotation and downstream analysis, we generated a gene activity matrix using Cicero, with a cell bin size of 80 and a minimum coaccessibility cutoff of 0.1^28^. We then correlated the average gene activities of each scTHS-seq cluster with the average gene expression of each snDropSeq cluster to identify rough cell types for each scTHS-seq cluster. scTHS-seq clusters that mapped to the same snDropSeq cluster were merged. We then validated scTHS-seq cell types by using Seurat to find gene activity markers for each scTHS-seq cell type, merging any cell types that lacked distinct markers. We further validated scTHS-seq cell types by correlating the average gene activities of each annotated scTHS-seq cell type with the average gene expression of each annotated snDropSeq cell type.

### HAR/HGE Annotation and Cell Type Enrichment

We compiled HARs, fetal HGEs (defined using fetal human and chimp brain datasets), Adult HGEs/HLEs (HGEs and HLEs defined using adult human and chimp brain datasets)^5–12^. We also sampled 20,000 ENCODE DNAse I accessible sites across the human genome to generate a set of control genomic regions^75, 76^. We annotated the genomic location of HARs and HGEs using ChIPSeeker, with promoters defined as being within 3 kb of a Transcription Start Site (TSS)^77^.

We computed a pseudobulk accessibility matrix by binarizing our peaks by cells matrix so that each peak is either accessible or not in any given cell and then summing the binary peak counts across all cells within a cell type, giving us a peaks-by-cell-types pseudobulk matrix. To identify cell types with a higher than expected accessibility of HARs/HGEs, we used a modified version of chromVAR^35^. Briefly, for each HAR/HGE type, we identified peaks that overlapped a HAR/HGE region and then used chromVAR to compute the deviation from expected accessibility for each peak. chromVAR then identifies a set of background peaks with similar GC content and average accessibility and computes the deviation for each background peak. chromVAR computes Z-scores by subtracting the peak deviation from the average background deviation and dividing by the standard deviation of the background deviation.

### HAR/HGE Transcription Factor Analysis

For each set of HARs/HGEs, we subsetted our peaks by cell types pseudobulk matrix to only those peaks overlapping that set of HARs/HGEs and then used chromVAR with JASPAR motifs to identify transcription factor motifs with cell type specific accessibility patterns^78^. Briefly, for each transcription factor motif, chromVAR identifies peaks containing that motif and then computes a deviation from expected accessibility for those peaks and a set of GC matched background peaks. chromVAR computes Z-scores by subtracting the average background deviation from the true deviation and dividing by the standard deviation of the background peaks. We only considered Z-scores from transcription factor motifs in a given cell type when that transcription factor was expressed in a specific fraction of cells in the scRNA-seq dataset dataset (expression defined as having at least one UMI in a given cell), setting the Z-score to zero if the transcription factor did not meet the expression threshold. We set the expression threshold at 0.05 for the THS analysis and at 0.025 for the ATAC analysis. The matched scRNA-seq data for the ATAC analysis was more sparse as it was generated using a combinatorial indexing method instead of a droplet based method^21^. For each set of HARs/HGEs, we visualized the cell type specific activity of transcription factor motifs with a Z-score greater than 4 in at least one cell type. A Z-score of 4 roughly corresponds to an unadjusted p-value of 0.000063, which after applying a Bonferroni correction for simultaneously testing 386 TF motifs, results in a reasonable adjusted p-value of approximately 0.025.

To validate our transcription factor motif activity, we used a set of differentially accessible peaks from a published study comparing human and chimp organoid accessibility^44^. We computed the log2 fold enrichment for JASPAR TF motifs in the peaks that showed higher accessibility in human organoids. We then correlated the log2 fold enrichment in human specific peaks with the cell type specific HAR/HGE transcription factor motif activity using a Pearson correlation. To avoid spurious correlations in cell types with low Z-scores, we only computed correlations for cell types with at least one TF Z-score greater than 4. Otherwise the correlation for that cell type was set to zero.

### Computing Average HAR/HGE Accessibility and Average Gene Expression

We computed the average scaled accessibility for each cell type by applying the TF-IDF transform to the binarized single cell peaks matrix using Signac, and then computing the average normalized accessibility for each cell type. To compute the average scaled gene expression for each cell type, we used Seurat’s NormalizeData to normalize the single cell gene expression counts and then used ScaleData to scale the normalized counts matrix^27^. We then took the average scaled expression for each cell type.

### Linking Transcription Factors to Genomic Regions (HARs/HGEs)

We first linked TFs to peaks overlapping HARs/HGEs by finding peaks that contained the TF binding motif which also overlapped one or more HARs/HGEs. Since the presence of a TF binding motif does not mean the TF is actually bound in a given cell type, we pruned the TF – peak links by correlating the overall TF motif enrichment with peak accessibility. Specifically, we first binned our single cell chromatin accessibility matrix into bins of 50 cells each using the binning method from the Cicero package^28^. Binning improves correlations with individual peaks since the single cell chromatin accessibility peaks matrices tend to be sparse^28^. We then ran our HAR/HGE TF motif enrichment on the binned peaks matrix using chromVAR, and correlated the motif enrichment Z-scores of each TF with the log(TF-IDF) transformed accessibility of all peaks with the TF motif present that also overlap an HAR/HGE. We pruned any TF – peak links with a Pearson correlation of less than 0.1. We used the dataset specific accessible peaks called via SnapATAC/MACS2 that overlap HARs/HGEs instead of using the HAR/HGE regions directly. HARs are defined by sequence acceleration while HGEs/HLEs are defined by the presence or absence of enhancer specific epigenomic marks. In both cases, the boundaries of these regions may not be well defined.

### Linking Genomic Regions (HARs/HGEs) to Genes with Cell Type Specific Hi-C Datasets

We used cell type specific chromatin conformation capture (Hi-C) data from a published study on the developing human cortex to link HARs/HGEs to genes by identifying peaks overlapping HARs/HGEs that are in physical contact with the promoter region (within 3kb of the TSS) of a gene^23^. We required that a given peak be accessible (average scaled accessibility greater than zero) in the given cell type in order to include it in the cell type’s regulatory network. We did not require that the contacted gene be expressed since we are not sure if the HAR/HGE is an activator or repressor.

### Linking Transcription Factors to Genes via HARs/HGEs

We then linked transcription factors (TFs) to genes by merging our previously generated TF – peak links and our cell type specific peak to gene links. We filter the TFs in each cell type specific network for TFs that are expressed (at least 5% of cells have non-zero expression of the TF in the THS dataset, 2.5% in the ATAC dataset).

### Fetal Human Gained Enhancer Network Visualization and Analysis

We visualized the TF regulatory networks for fetal HGEs using force directed layouts with the R igraph package^79^. We highlighted the TFs with a fetal HGE motif enrichment Z-score of greater than 4 within each network. We computed the undirected Eigenvector Centrality for all nodes in each network using the eigen_centrality function in igraph with default parameters. We also computed the undirected Edge Betweenness Centrality for all edges in each network using the edge_betweenness function.

### NFIC and LHX2 network analysis

We used Fisher’s exact test and the MSigDB Hallmark Genesets and Canonical Pathway Genesets to compute pathway enrichment for NFIC target genes in the **RG** and **ipEx** networks^47, 48^. We compared the pathway enrichment of the scTHS and sciATAC networks using heatmaps the –log(p-values) for the top 5 enriched pathways for each. We then identified NFIC consensus target genes present in both the sciATAC and scTHS cell type networks and built consensus sub-networks using those genes as well as any other TFs targeting those genes. We merged the **RG** and **ipEx** sub-networks into a single network for visualization, highlighting TFs in red. We applied the same target gene analysis to the LHX2 target genes, building a consensus sub-network of all LHX2 target genes across the **RG** and **ipEx** networks along with any TFs also targeting those genes.

We visualized the Hi-C contacts between fetal HGEs with NFIC and LHX2 motifs and key target genes (GLI2 and C1orf61 respectively) using the merged **RG** and **ipEx** HGE – gene networks from the scTHS-seq dataset. We added the accessibility tracks for **RG**, **ipEx**, and mature **Ex** cell types by generating bigwig files from all cells in each respective cell type using the bamCoverage command with the parameters “-bs 50 –normalizeUsing CPM –skipNAs”. Accessibility tracks were generated from the scTHS dataset, and the **RG** cells included both **oRG** and **vRG** cells while the mature **Ex** included all layer specific neurons (**ExL2/3, ExL4, ExL5/6**) We generated the initial plot using the plot_connections function from Cicero and added the accessibility tracks using the Gviz package^28, 80^.

### Fetal Human Chromatin Atlas Analysis

We downloaded the cerebrum chromatin accessibility (generated using scATAC-seq) (https://atlas.brotmanbaty.org/bbi/human-chromatin-during-development/) and gene expression datasets (https://atlas.brotmanbaty.org/bbi/human-gene-expression-during-development/) from their respective fetal Human Atlases^20, 21^. Due to the extremely large size of the RNA dataset (over 2 million cells), we down-sampled the data to 10,000 cells per cell types. For both the chromatin and RNA datasets, we reclustered the Excitatory (Ex) neuron clusters using the pre-computed UMAP coordinates from the databases to generate finer resolution clusters. We correlated the average scaled gene activity or gene expression from the chromatin and RNA datasets with the average gene activity or average gene expression from our scTHS-seq/snDrop-seq datasets. We also visualized the average gene activities of canonical Radial Glia (*GLI3, VIM, NES, FABP7, SOX2*), Intermediate Progenitor (*EOMES, PPP1R17, NEUROG2, PAX6*), and Astrocyte (*S100B, NEU1*) marker genes in the scATAC-seq cell types.

We then re-mapped the cell types using the marker gene activity/expression and the average activity/expression correlations. For the sciATAC chromatin dataset, given that the astrocyte cell type expressed radial glia marker genes at a higher level than astrocyte markers and was more highly correlated with radial glia gene activity than astrocyte activity, we relabeled the scATAC Astrocytes as radial glia (**RG**). We identified that Cerebrum Unknown Cell Type 3 was most likely intermediate progenitors (**ipEx**) and relabeled those cells accordingly. We also found 2 Excitatory Neuron sub-clusters that mapped well to **ebEx** and relabeled those sub-clusters. The remaining excitatory neuron sub-clusters were labeled **Ex**. For the sciRNA dataset, we also relabeled astrocytes as **RG** cells. We also identified Ex sub-clusters in the sciRNA dataset that corresponded to **RG**, **ipEx**, and **ebEx** and re-labeled those sub-clusters accordingly. We visualized the relabeled cells using the pre-computed UMAP coordinates.

We also lifted over the scATAC-seq dataset peaks (which were mapped to the hg19 reference build) to the hg38 reference build.

## Supplemental Figures

**Figure S1:**
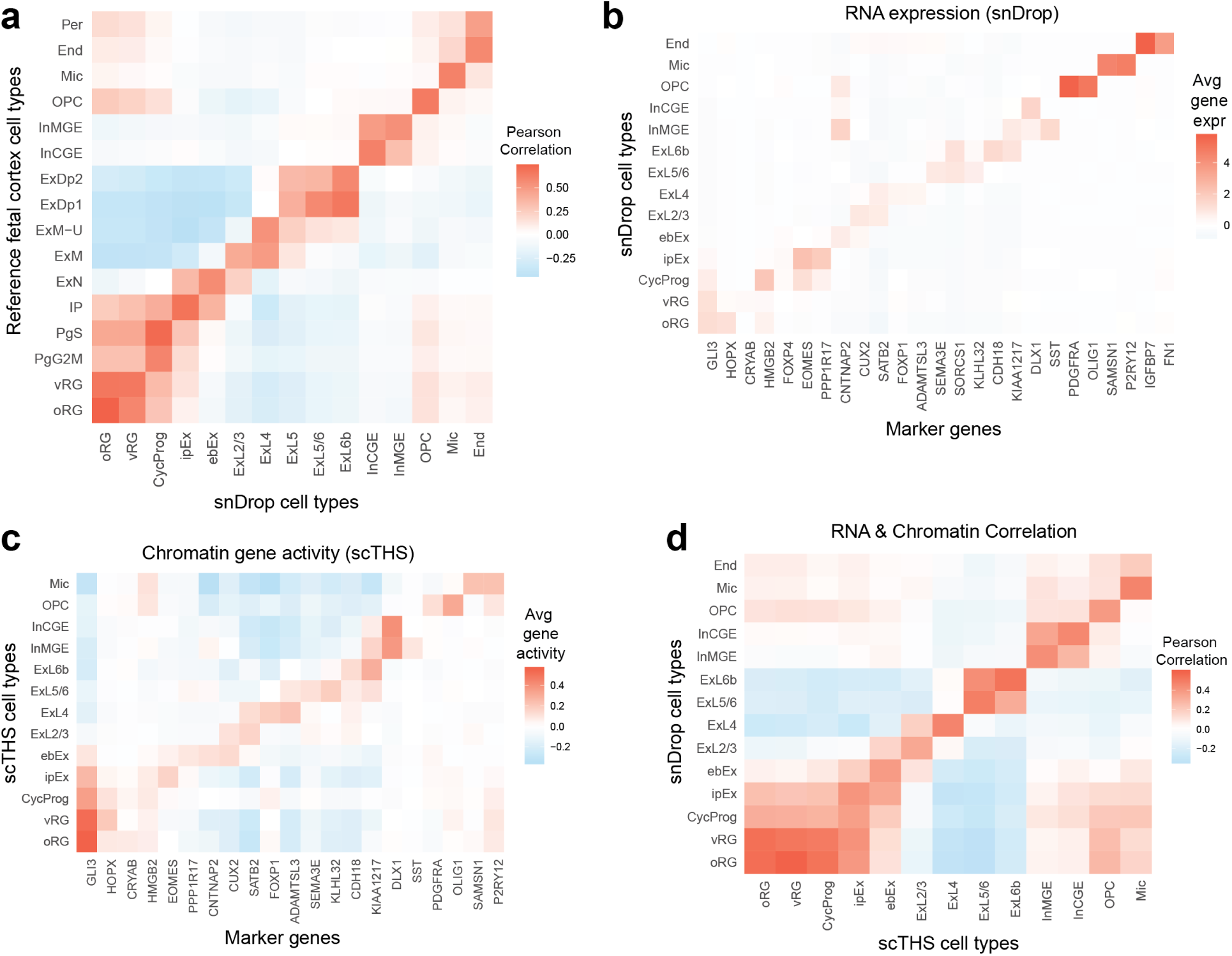
**a)** Pearson correlation of average cell type gene expression between cell types from a reference developmental cortex dataset and cell types from our fetal snDrop-seq dataset. **c)** snDrop-seq gene expression heatmap of key cell type markers. **b)** scTHS-seq chromatin gene activity heatmap of key cell type markers. **d)** Pearson correlation between average snDrop-seq cell type gene expression and average scTHS-seq chromatin gene activity.

**Figure S2:**
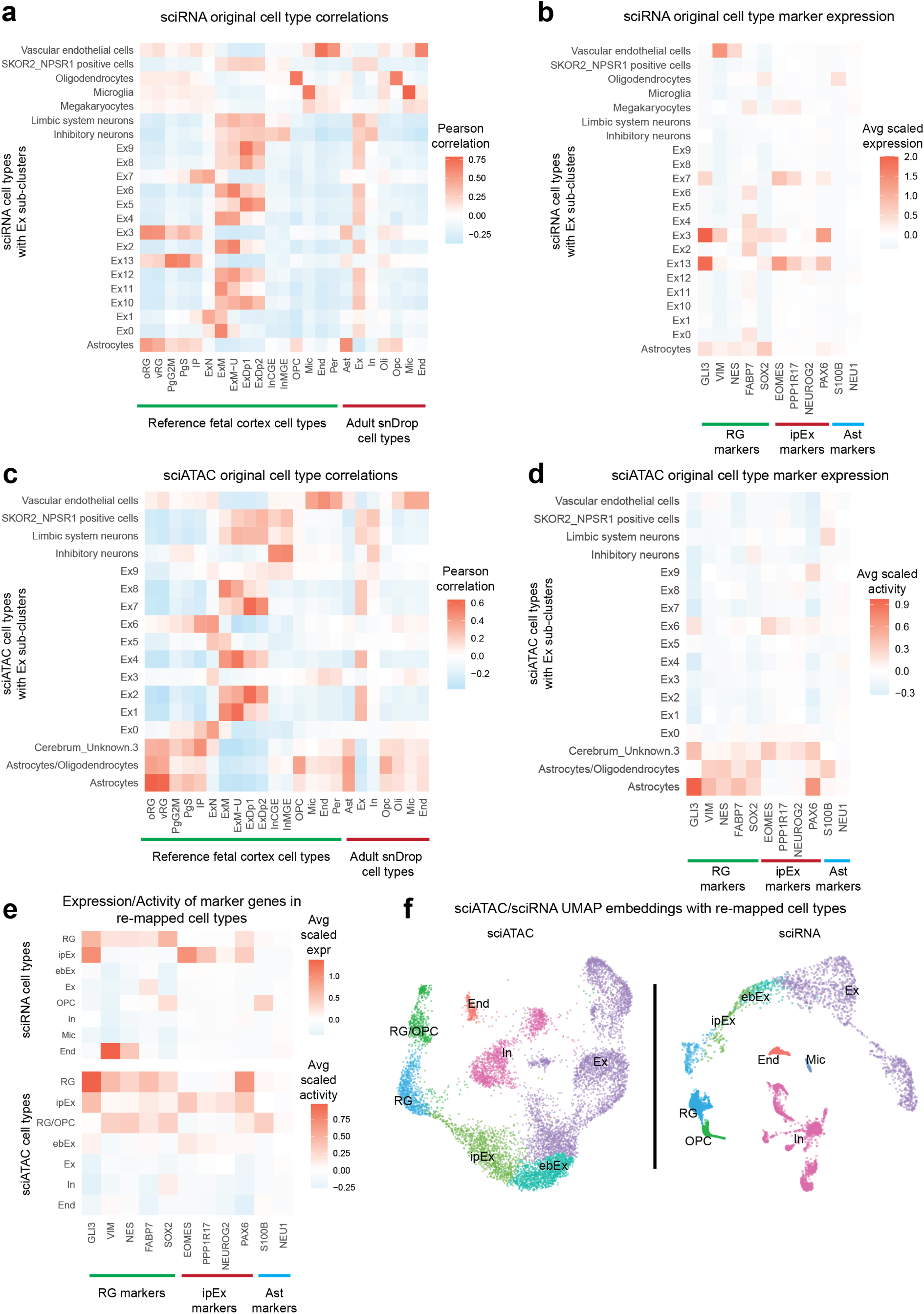
**a)** Heatmap of Pearson correlations between the average scaled gene expression of the original sciRNA cell types (with **Ex** sub-clusters) and reference developmental cortex RNA cell types as well as our adult snDrop cell types. **b)** Average scaled expression of **RG**, **ipEx**, and **Ast** markers in the original sciRNA cell types. **c)** Heatmap of Pearson correlations between the average scaled gene activity of the original sciATAC cell types (with **Ex** sub-clusters) and reference developmental cortex RNA cell types as well as our adult snDrop cell types. **d)** Average scaled gene activity of **RG**, **ipEx** and **Ast** markers in the original sciATAC cell types. **e)** Average scaled gene expression **RG**, **ipEx** and **Ast** markers in the re-mapped sciRNA and sciATAC cell types. **g)** UMAP embeddings of the sciATAC and sciRNA datasets with re-mapped cell types.

**Figure S3:**
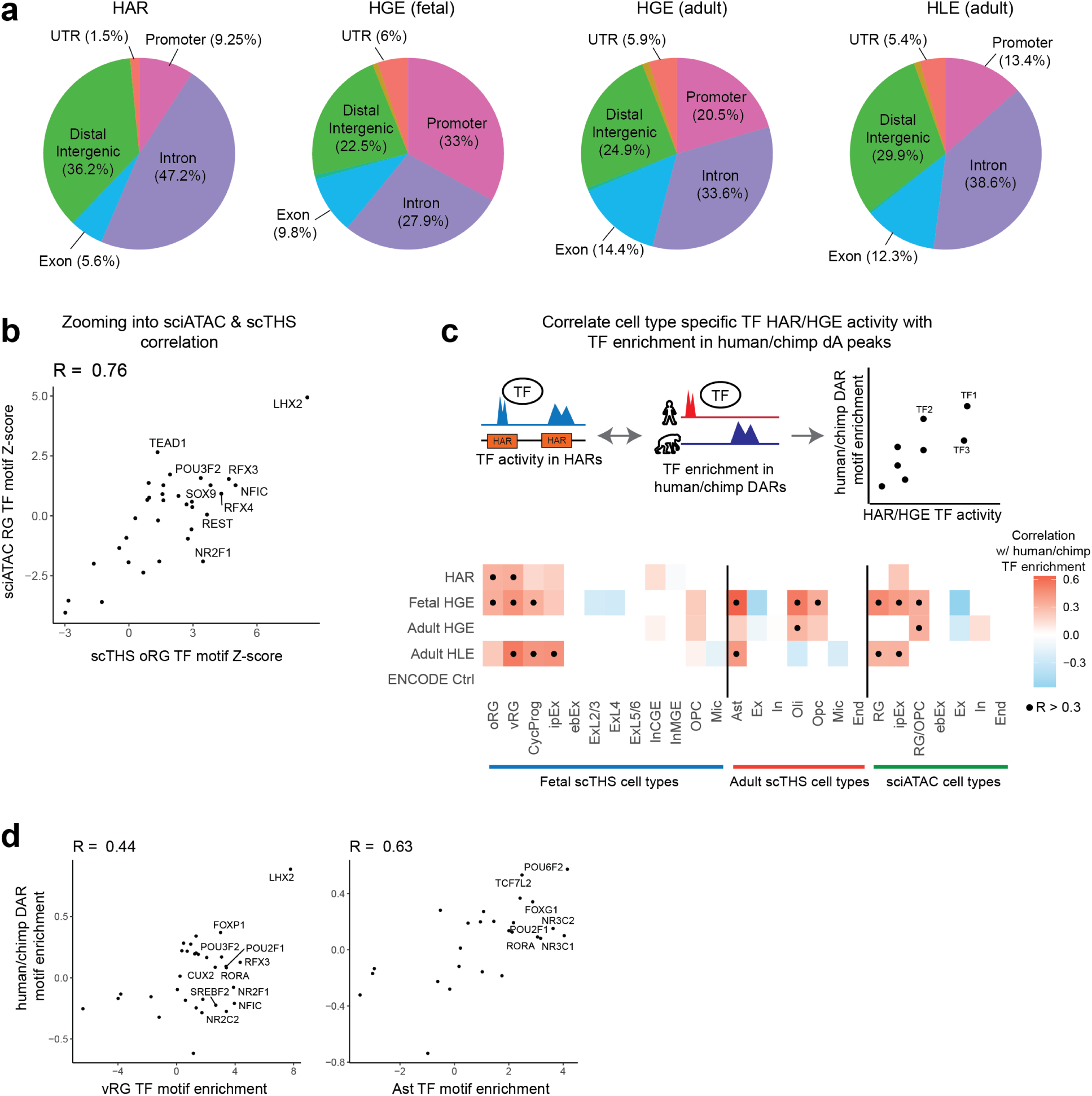
**a)** Genomic annotation of HAR/HGE regions. Promoters are defined as regions within 3kb of a transcription start site (TSS). **b)** Scatterplot of Pearson correlation of TF motif enrichment in fetal HGEs between our scTHS **oRG** cell type and the sciATAC **RG** cell type. **c)** Heatmap of Pearson correlations between TF motif enrichment in HARs/HGEs from scTHS and sciATAC cell types and TF motif enrichment in regions that are differentially accessible (DARs) between human and chimp brain organoids. **d)** Scatterplots comparing TF motif enrichment in fetal HGEs for scTHS **oRG**/**Ast** and TF motif in enrichment in human/chimp DARs.

**Figure S4:**
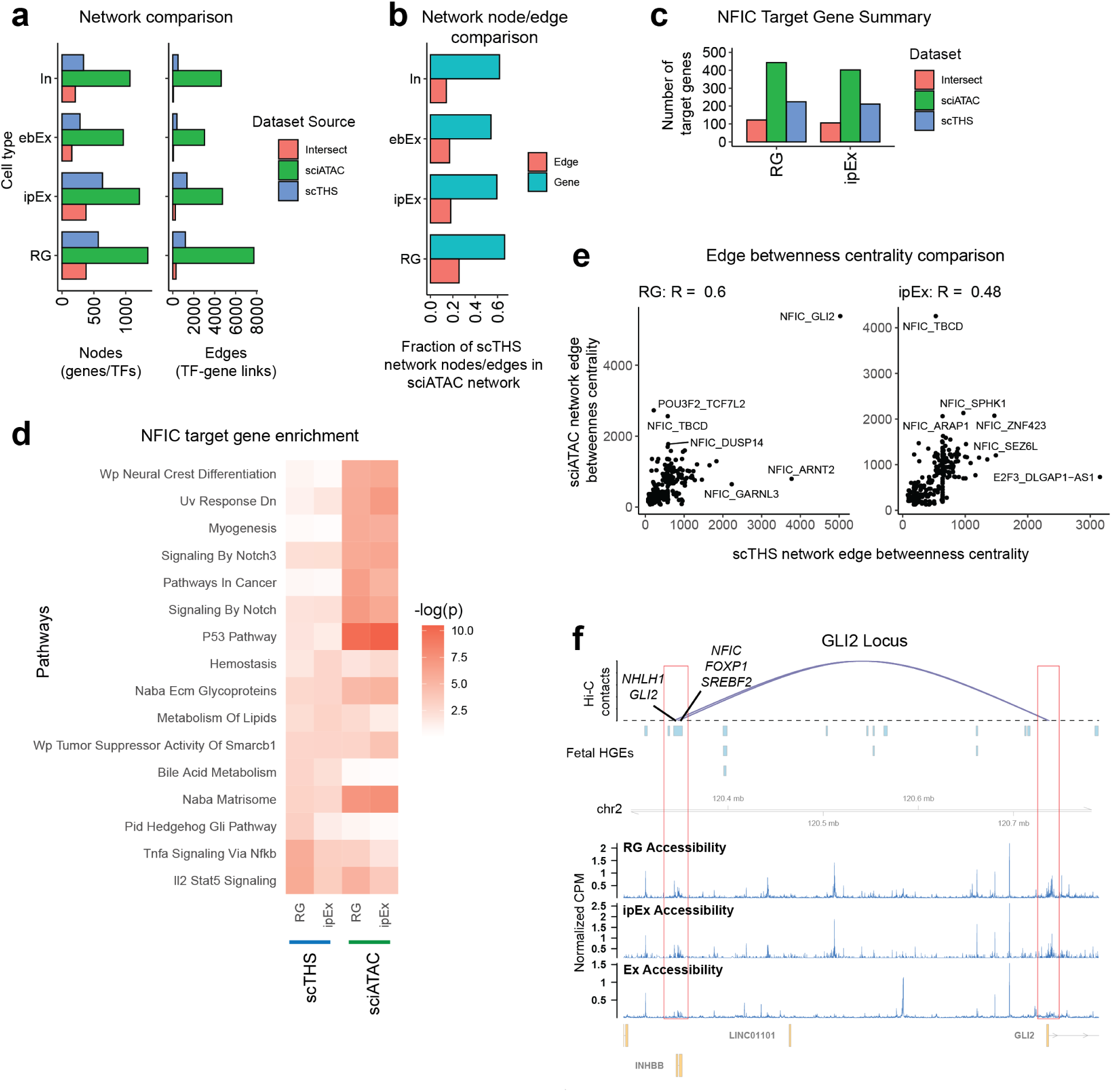
**a)** Bar plot comparing the number of nodes (genes & TFs) in each fetal HGE network for sciATAC and scTHS datasets as well as the number of nodes shared between corresponding networks from each dataset. **d)** Bar plot showing the fraction of edges/nodes present in scTHS networks that are also found in corresponding sciATAC networks. **c)** Bar plot comparing the number of *NFIC* target genes in sciATAC and scTHS fetal HGE TF networks. **d)** Heatmap visualizing the top pathways enriched for NFIC targets across sciATAC and scTHS cell type networks. **e)** Scatterplots visualizing the influence of edges (TF – gene) links by plotting edge betweenness centrality across the sciATAC and scTHS networks. Edge betweenness centrality scores how often each edge serves as the shortest path between different parts of the network, thus enabling us to identify the most influential TF – gene links. **f)** Visualization of the fetal HGEs with NFIC/FOXP1/SREBF2 and NHLH1/GLI2 motifs that contact the C1orf61 promoter. Arcs represent a chromatin contact from neural progenitor (**RG/ipEx**) Hi-C networks. Tracks showing the normalized accessibility of the locus in Counts per Million (CPM) are displayed for **RG**, **ipEx** and **Ex** cell types.

